# Increased Beta-Band Input to Spinal Motor Neurons is Not Confined to the Responding Muscle During a Go/No-Go Task

**DOI:** 10.64898/2026.05.16.725172

**Authors:** François Hug, François Dernoncourt, Clément Naveilhan, Wolbert Van den Hoorn

**Author notes:** Corresponding author Prof. François Hug Université Côte d’Azur, LAMHESS, Nice, France.

## Abstract

During movement preparation and inhibition, the sensorimotor cortex shows alpha-and beta-band oscillations that are also expressed in the discharge of spinal motor neurons. Whether this oscillatory activity is confined to the motor neurons innervating the muscles involved in the prepared action is unknown. Here we combined electroencephalography with recordings of spinal motor neurons innervating the tibialis anterior. Participants held a submaximal dorsiflexion while performing a Go/No-Go task in which the instructed response was either a ballistic dorsiflexion or handgrip, making the tibialis anterior the responding or a non-responding muscle, respectively. Alpha-and beta-band power increased after the imperative cue in both cortical activity and motor neuron output, including when the tibialis anterior was not involved in the response. Because power modulations in motor neuron output can be confounded by the concurrent change in discharge rate, we assessed the oscillatory activity shared across motor neurons by calculating coherence between disjoint subsets, which is largely insensitive to discharge rate. We then used simulation-based inference to identify synaptic input modulations compatible with our observations. These analyses indicated that beta-band common input increased whether or not the tibialis anterior was involved in the response, whereas alpha-band input increased only when it was the responding muscle. The concurrent decrease in discharge rate and torque was attributed to reduced net excitatory drive, whereas alpha-and beta-band inputs accounted for the increase in synchronisation. These results demonstrate that cue-related oscillatory inputs reach spinal motor neurons more broadly than classically assumed, and shape motor neuron synchronisation.

**Key Points:**

- Alpha-and beta-band cortical oscillations accompany movement preparation and inhibition, but whether they are expressed in muscles not involved in the response is unknown.
- We combined electroencephalography with recordings of tibialis anterior motor neurons during a Go/No-Go task in which the instructed response was either a dorsiflexion or a handgrip.
- Beta-band common input to these motor neurons increased after the cue even when the handgrip was the instructed response, whereas the increase in alpha-band input occurred only when the dorsiflexion was the instructed response.
- Motor neuron synchronisation increased in both tasks, and modelling attributed it to this increase in this oscillatory activity.

## Introduction

Brain oscillations reflect the synchronized activity of neuronal populations and serve as a fundamental mechanism for communication and information processing across cortical networks (Buzsáki & Draguhn, 2004). Oscillations in the alpha (8–12 Hz) and beta (13–30 Hz) bands have been linked to distinct aspects of motor control, including movement preparation, sensorimotor integration, response inhibition, and the control of ongoing motor output (Klimesch, 2012; Picazio *et al*., 2014; Little *et al*., 2019; Wessel, 2020). The peripheral expression of these oscillatory signals in spinal motor neuron output is less well documented. In particular, it remains unclear whether alpha-and beta-band oscillations selectively reach the motor pools directly engaged in a motor task or are expressed more broadly across the motor system. This distinction is functionally important: selective routing would support effector-specific control, whereas broader expression would be consistent with either a general state-setting signal not restricted to a single effector, or an incidental leak of central oscillatory activity.

Go/No-Go and stop-signal tasks, in which participants must either execute a prepared movement or rapidly inhibit it in response to an imperative cue (Picazio *et al*., 2014; Zicher *et al*., 2024), are useful paradigms for experimentally inducing well-characterized modulations of alpha-and beta-band activity. Specifically, successful movement inhibition is marked by increases in alpha-and beta-band power following the cue (Pani *et al*., 2014; Zicher *et al*., 2024). Post-cue beta-band modulations have also been observed in spinal motor neuron output during No-Go conditions, but so far only in the muscle directly involved in response selection (Zicher *et al*., 2024), consistent with previous observations of beta-band cortico-muscular coupling during isometric contractions (Baker, 2007).

Empirical evidence for broad expression of alpha-and beta-band oscillations across motor neuron pools remains limited, as most studies have focused on the primary effector (Zicher *et al*., 2024) or have relied on indirect measures such as surface electromyography (EMG) or force (Rangel *et al*., 2024; De Havas *et al*., 2025). Using transcranial magnetic stimulation, studies have shown that changes in corticospinal excitability during movement preparation and inhibition can extend beyond the selected effector to task-irrelevant muscles (Duque *et al*., 2010; Greenhouse *et al*., 2015). Similarly, stopping a motor response can be accompanied by a transient reduction in force or muscle activation in effectors not involved in that response (Rangel *et al*., 2024; De Havas *et al*., 2025). However, these measures do not reveal whether task-related oscillatory dynamics are similarly broadly expressed across motor pools. Without direct measures of the neural drive, it is impossible to determine whether these behavioural changes reflect the propagation of alpha-and beta-band oscillations themselves or instead reflect changes in the low-frequency neural drive that directly modulates muscle force (Farina *et al*., 2016), potentially through decreased corticospinal excitability (Leocani *et al*., 2000). Resolving this is essential not only for understanding the bandwidth and specificity of muscle control, but also for determining whether high-frequency oscillations measured at the muscle level can serve as an independent control signal for neural interfacing and movement augmentation (Ibanez *et al*., 2025).

Because spinal motor neurons constitute the “final common pathway” (Sherrington, 1906) through which descending and peripheral inputs are integrated, recording their activity provides a direct way to determine whether cortical alpha-and beta-band oscillations are expressed at the peripheral level. Crucially, detecting these oscillatory signals requires a sufficient number of spikes over time, which in practice mainly depends on sampling a large number of motor neurons whose collective output can reliably capture high-frequency dynamics (Negro & Farina, 2011). In this study, we combined large-scale motor neuron recordings (up to 57 motor neurons per participant) with electroencephalography (EEG). Participants performed Go/No-Go tasks while holding a submaximal dorsiflexion. The instructed response was either a ballistic dorsiflexion or a ballistic handgrip, making the recorded muscle (tibialis anterior, TA) either the responding or a non-responding muscle, respectively. We aimed to determine whether alpha-and beta-band oscillations are expressed beyond the responding muscle and whether these peripheral oscillations have measurable consequences for motor output. We used simulation-based inference to identify the patterns of synaptic inputs most likely to account for the observed motor neuron output (Cranmer *et al*., 2020; Dernoncourt *et al*., 2026).

## Methods

### AI disclosure and data availability

We used ChatGPT (GPT-5.3, OpenAI) and Claude (Opus 5, Anthropic) to assist with language editing to improve clarity and readability, and to support code development for data analysis. All AI-assisted outputs were critically reviewed, edited, and verified.

### Participants

Eighteen individuals volunteered to participate in the experiment. None reported a history of upper or lower limb injury in the six months preceding the experiment. The study procedures were approved by a national ethics committee (CPP Sud-Méditerranée IV, n°2025-A02574-45) and were conducted in accordance with the declaration of Helsinki, except for registration in a database. All participants were fully informed of the potential risks and discomforts associated before providing written informed consent.

Due to technical issues, EMG data were not streamed together with EEG data in one participant, and reliable motor units could not be identified in two participants. Data are therefore reported for 15 participants (3 females, age: 32.2 ± 8.5 years, height: 175.4 ± 6.8 cm, body mass: 70.0 ± 7.7 kg).

### Procedures

The experimental session was based on Go/No-Go tasks known to elicit specific modulations of cortical oscillations. Participants were seated in a custom-built chair connected to an ankle ergometer (DinamometroGC, OT Bioelettronica, Torino, Italy). The right foot was securely attached to the footplate, with the ankle positioned at approximately 10° of plantarflexion (Fig. 1). The knee was flexed at 20° and the hip at 80° (0° corresponding to full extension). Participants also held a handgrip dynamometer (Cor2, OT Bioelettronica, Torino, Italy) in their right hand.

**Figure 1.**
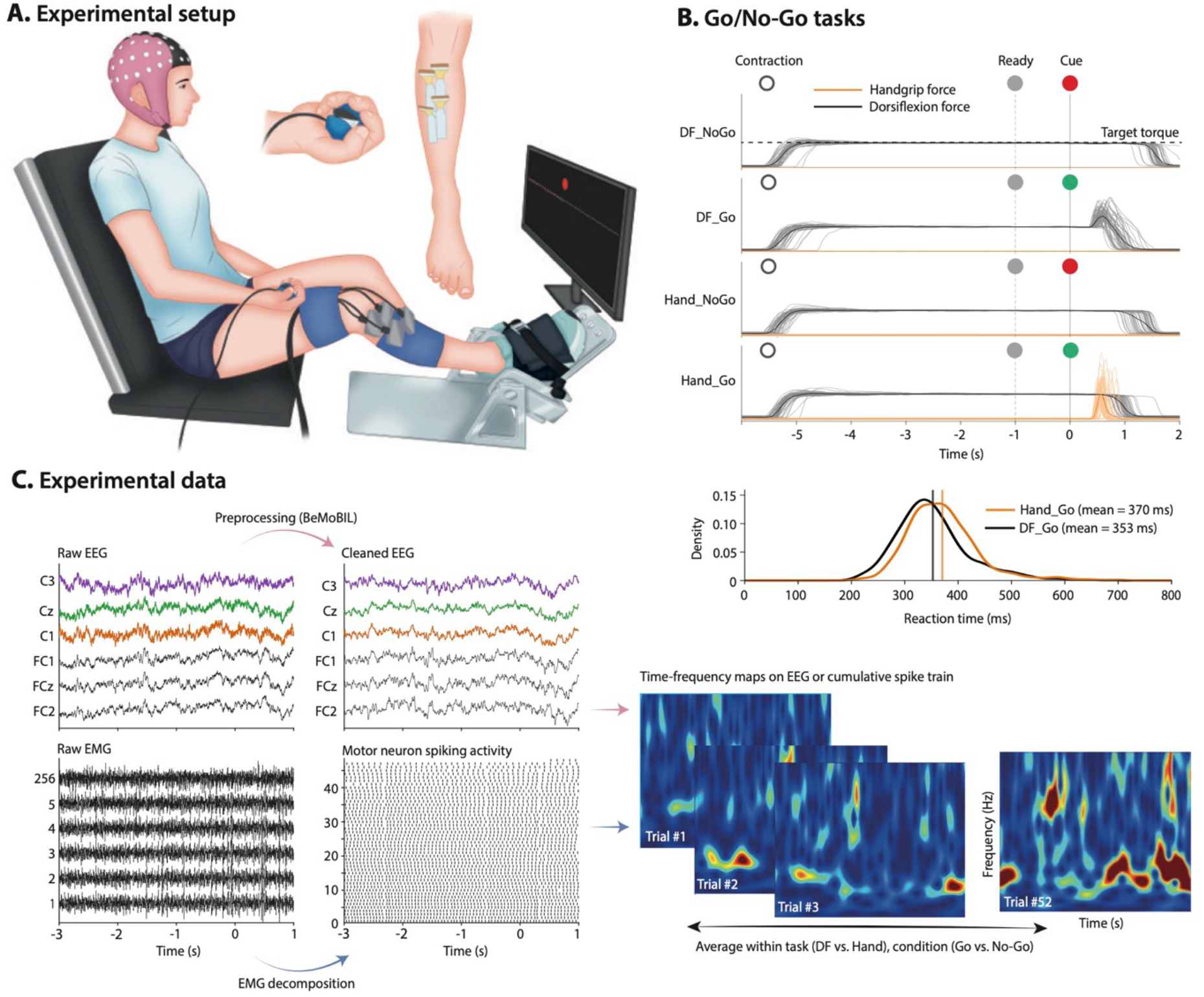
Experimental design. **(A)** Participants performed isometric dorsiflexion tasks while seated with the right foot secured to an ankle ergometer and the right hand holding a handgrip dynamometer. High-density surface electrode grids (256 channels in total) were placed over the tibialis anterior, and a 64-channel electroencephalography (EEG) cap recorded brain activity. **(B)** Each trial began with an instruction cue (open circle) indicating that participants should initiate dorsiflexion at submaximal target torque. After 5 s, a ready cue (grey filled circle) was presented, followed 1 s later by an imperative cue indicating either a Go (green) or No-Go (red) trial. In DF_Go trials, participants performed a ballistic dorsiflexion. In Hand_Go trials, participants performed a ballistic handgrip contraction while maintaining dorsiflexion. In DF_NoGo and Hand_NoGo trials, participants were instructed to maintain the ongoing dorsiflexion contraction without producing an additional response. The bottom panel shows the reaction time distributions across participants for DF_Go (black) and Hand_Go (orange). **(C)** Electromyography (EMG) and EEG signals were recorded simultaneously. EMG signals were decomposed into motor neuron spike trains. For each trial, time-frequency maps were computed from cleaned EEG and from the cumulative spike train and then averaged across trials to quantify alpha-and beta-band modulations. DF, dorsiflexion.

The session began with a standardized warm-up, followed by two maximal voluntary dorsiflexion contractions and two maximal handgrip contractions, performed in an interleaved order. Each contraction lasted ∼3 s, with 60 s of rest between contractions. Participants then performed three task blocks in a randomized order: five 20-s submaximal isometric contractions at 12.5 Nm, a Go/No-Go task involving dorsiflexion, and a Go/No-Go task involving handgrip. Each Go/No-Go task was divided into 4 blocks of 26 trials (13 Go, 13 No-Go), yielding 8 blocks in total (4 dorsiflexion, 4 handgrip), and 52 No-Go trials per task. The relatively large proportion of No-Go trials, compared with typical Go/No-Go paradigms, was motivated by the need to obtain a sufficient number of No-Go trials while limiting fatigue from sustained dorsiflexion contractions. This was particularly important as No-Go trials were the primary condition of interest due to their well-characterized modulation of beta-band activity (Zicher *et al*., 2024).

Participants received visual cues displayed on a screen together with the torque target (Fig. 1). First, an instruction cue (open circle), accompanied by the display of the target torque, indicated that participants should produce dorsiflexion at the target torque (12.5 Nm, i.e. 32±11% MVC). After 5 s, a ready cue (grey filled circle) was presented, followed 1 s later by an imperative cue indicating the trial type: a green filled circle for Go trials and a red filled circle for No-Go trials. During Go trials in the dorsiflexion task (DF_Go), participants were instructed to produce a rapid ballistic dorsiflexion above the maintained torque, as quickly as possible following the imperative cue. During Go trials in the handgrip task (Hand_Go), participants were instructed to squeeze the dynamometer as quickly as possible while maintaining dorsiflexion at the submaximal target torque (12.5 Nm). During No-Go trials, participants were instructed to maintain dorsiflexion at the target torque until the imperative cue disappeared, 1 s after onset. Trials were separated by 7 s on average (jittered by ±20%), and each block lasted approximately 6min30s.

Following visual inspection, trials with incorrect responses (missed responses to Go cues, or responses to No-Go cues) were excluded, corresponding to an average of 3.0 ± 2.1 trials per participant (DF_NoGo: 6.7 ± 6.0, Hand_NoGo: 3.4 ± 2.9, DF_Go: 0.5 ± 1.1, Hand_Go: 0.8 ± 0.9).

### Electroencephalography recordings and processing

EEG data were collected using a 64-channel Waveguard Original cap equipped with gel-based Ag/AgCl electrodes (ANT Neuro, Germany). Signals were amplified and digitized at 500 Hz using an eego mylab system (ANT Neuro, Germany). CPz and AFz served as the reference and ground electrodes, respectively. During preparation, electrode impedances were reduced to below 10 kΩ, and signal quality was visually inspected to ensure absence of excessive noise before data acquisition.

EEG data were pre-processed using EEGLAB (Delorme & Makeig, 2004) functions (v. 2025.0.0) and the BeMoBIL pipeline (Klug *et al*., 2022), following the procedure described in Naveilhan et al (2025). Preprocessing was applied to the concatenated EEG data from all eight Go/No-Go blocks and the trapezoidal contraction task. First, spectral artifacts were attenuated using Zapline-plus (Klug & Kloosterman, 2022), and noisy channels were identified using the BeMoBIL processing routine, removed, and subsequently interpolated using spherical splines. The data were then re-referenced to the common average across scalp electrodes, excluding the EOG channel. Before running adaptive mixture independent component analysis (AMICA), a temporary high-pass filter was applied (lower edge = 2 Hz, filter order = 1650; corresponding to an effective cutoff of approximately 1.5 Hz). AMICA automatic sample rejection was enabled (10 rejection iterations, sigma threshold = 3). Independent components were classified into seven categories (brain, muscle, eye, heart, line noise, channel noise, and other) using ICLabel with the “lite” setting (Pion-Tonachini *et al*., 2019). Components were retained if they were classified as ≥ 50% brain-related and exhibited a residual variance below 15%. Using these criteria, a conservative average of 11.9 ± 5.0 components per participant were retained (range: 4-21). Component weights, equivalent dipoles, and labels were then copied to the unfiltered data, and the retained components were filtered between 0.5 and 50 Hz and projected back to electrode space to reconstruct the cleaned EEG signal for each channel.

### Surface electromyography recordings and decomposition

EMG signals were recorded from the TA muscle using four adhesive grids of 64 electrodes (256 electrodes in total; 4-mm inter-electrode distance, HD04MM1305, OT Bioelettronica, Italy; Fig. 1). Prior to electrode placement, the skin was shaved and cleaned with an abrasive paste (Nuprep, Weaver and Company, USA). The grids were secured with double-sided adhesive foam layers (SpesMedica, Battipaglia, Italy), and skin-electrode contact was established by filling the cavities of the adhesive layers with conductive paste (SpesMedica, Battipaglia, Italy). A reference electrode (5 × 5 cm; Kendall Medi-Trace; Covidien, Ireland) was placed over the tibia, and a water-damped strap electrode (ground electrode) was placed around the left ankle. EMG signals were recorded in a monopolar montage, bandpass filtered (10–500 Hz), and digitized with 16-bit precision at a sampling rate of 2000 Hz using a multichannel acquisition system (Novecento+; 960-channel EMG amplifier; OT Bioelettronica, Italy).

EMG signals were decomposed using convolutive blind source separation (Negro *et al*., 2016a), as implemented in the MUedit software (Avrillon *et al*., 2024). Prior to automatic decomposition, all channels were visually inspected, and those with a low signal-to-noise ratio or visible artifacts were discarded. The resulting spike trains were manually edited according to established procedures (Del Vecchio *et al*., 2020). Manual editing consisted of removing detected peaks that resulted in erroneous firing rates (outliers) and adding missed firing times that were clearly distinguishable from the noise. Motor unit pulse trains were then recalculated with updated separation vectors and accepted once all putative firings were selected. This procedure has been shown to be highly reliable across operators (Hug *et al*., 2021).

Decomposition was performed independently for each electrode grid. Because the same motor units could be identified across multiple grids covering the same muscle, duplicate units were detected by comparing their spike trains. Firings occurring within a 0.5-ms interval across two spike trains were considered common; and motor units sharing more than 30% of their firings were considered as duplicates. When duplicates were identified, only the motor unit with the lowest coefficient of variation of inter-spike intervals was retained.

### Synchronisation of electroencephalography and electromyography signals

EEG and EMG signals were synchronized using LabRecorder via the Lab Streaming Layer (LSL) protocol (Kothe *et al*., 2025). Both EEG and EMG acquisition systems streamed data in real time to LSL, providing a common time reference across modalities. LabRecorder stored these streams with precise timestamps, allowing accurate alignment for offline analysis. This approach compensates for device-specific latencies and clock differences, providing sub-millisecond synchronisation accuracy between EEG and EMG signals.

### Time-frequency analysis

Time-frequency maps were computed separately for EEG and cumulative spike trains (CST) using the superlet transform (Moca *et al*., 2021). EEG data were epoched from-4 to 3 s relative to the imperative cue (Go or No-Go) and CST data from-5 to 2 s, in both cases with buffers of 1–2 s on either side of the −3 to 1 s window retained for analysis, so that edge effects fell outside it. For EEG, time-frequency power was computed for each EEG channel, and then averaged across Cz, C1, and C3 for the main analyses. For CST, binary motor unit spike trains were summed, and the resulting signal was demeaned and detrended. Time-frequency decomposition was performed from 4 to 40 Hz in 0.5-Hz steps, with a time step of 10 ms for EEG and 4 ms for CST. Both used a multiplicative superlet combination with a base width of 2 cycles, a Gaussian width of 3, and an adaptive order increasing with frequency from 4 to 12.

Time-frequency maps were normalized separately for each task. Specifically, normalization was performed trial-wise using a two-step z-scoring procedure (Grandchamp & Delorme, 2011). First, raw power at each frequency was z-scored within each trial over the full analysis window. Second, these values were re-normalized using the mean and standard deviation computed from the pooled reference period across trials (-3 to-2 s). The resulting normalized trial maps were averaged across kept trials within each task to obtain participant-level time-frequency maps.

### Coherence between motor unit spike trains

Because motor units discharge quasi-regularly, the CST power spectrum shows a peak at the mean discharge rate, which was close to the upper edge of the alpha band in our data. A decrease in discharge rate could therefore shift part of this peak into the alpha band (8–12 Hz) and increase alpha power, with no change in alpha-band input. To determine whether the changes in alpha-and beta-band power reflected changes in oscillatory activity shared across motor neurons, we calculated the coherence between two cumulative spike trains, each built from a separate subset of motor units. As each motor unit belonged to only one subset, the cross-spectrum between the two subsets contained only terms from pairs of different units rather than terms reflecting the discharge rate of individual units. This makes coherence much less sensitive than CST power to changes in discharge rate (Negro *et al*., 2016b).

We calculated the pooled coherence on two CSTs computed from equally sized groups of motor units. Because motor units were identified separately in each run, the groups were formed within each run. The number of motor units in each group was the maximum possible, that is, half of the motor units identified in that run (on average 18 per group; range, 7–28). For each run, 100 random splits of the motor units into disjoint groups were drawn. We calculated the coherence using Welch’s periodogram with non-overlapping Hann windows of 1 s, each window corresponding to one trial. The same three windows as those used for the time-frequency maps were analysed, i.e., Baseline (−2 to −1 s), Ready (−1 to 0 s), and Post-Cue (0 to 1 s). The spectra were pooled across all trials of the four runs, and coherence was averaged across permutations. Coherence values were transformed using Fisher’s transformation (z’ = atanh √COH) and averaged within the delta (1–5 Hz), alpha (8–12 Hz) and beta (13–30 Hz) bands. The same trials and the same motor units were used in the three windows, and the analyses focused on changes in coherence between windows within each condition.

### Simulation-based inference

We used simulation-based inference [sbi Python toolbox (Tejero-Cantero *et al*., 2020)] to identify synaptic inputs compatible with the experimentally observed motor neuron behaviour. Inference was performed in two stages. First, we estimated baseline motor neuron pool properties and baseline input parameters. Second, conditioning on the posteriors from this first stage, we inferred cue-related modulations of net excitatory drive and alpha-and beta-band inputs.

Motor-neuron simulation model and input parameterization

The model consisted of a pool of 40 leaky integrate-and-fire neurons with size-dependent membrane and afterhyperpolarization properties (Caillet *et al*., 2022), implemented in Brian2 (Stimberg *et al*., 2019), and adapted from Dernoncourt et al. (2026). Model parameters and their fixed values or prior ranges are reported in Table S1, and we have made the code publicly available (see Data Availability).

We simulated spike trains over 10-s trials, with a 6-s analysis window (2 s trimmed on either side) to minimize edge artifacts and ensure that the model had reached a steady firing regime. Time was referenced to the imperative cue, as in the experimental Go/No-Go task; the analysis window spanned-4 to +2 s. Each motor neuron received a tonic excitatory drive, common fluctuating inputs, and independent noise. The common fluctuating inputs consisted of three zero-mean band-limited Gaussian processes: a low-frequency input (0–5 Hz), an alpha-band input (8–13 Hz), and a beta-band input (13–30 Hz). To reproduce the temporal dynamics of alpha and beta activity, alpha-and beta-band fluctuations were multiplied by non-negative modulating envelopes. Envelopes were generated as low-pass filtered Gaussian processes with cutoffs of 3 Hz and 10 Hz for alpha and beta, respectively, chosen to cover most (>80%) of the power in the experimental alpha-and beta-band Hilbert envelopes. Each common input component was scaled so that its standard deviation matched the corresponding amplitude parameter. Each motor neuron was assigned two scaling weights: one for the common input fluctuations and one for the independent noise, with configurable distributions of those weights across neurons.

Cue-related modulations of the net excitatory drive and the alpha-and beta-band inputs were modelled as transient bursts plus linear trends, yielding six cue-related parameters in total (one burst amplitude and one trend slope per input component). For the net excitatory drive, the burst and trend were added directly to the tonic baseline. For the alpha-and beta-band inputs, they were added to the modulating envelopes, which were clipped at zero to maintain non-negative amplitudes. Bursts were Gaussian-shaped (centre: 0.25 s post-cue; SD: 100 ms) and trends were linear, beginning at-1 s and extending to the end of the simulation.

Observable features

The same features were extracted from simulated and experimental data. The CST was obtained by summing spike trains across active motor neurons and normalizing by the number of active neurons. Four time-varying profiles were then derived: low-frequency, alpha-band, and beta-band CST modulations, together with a motor neuron synchronisation index. The low-frequency CST modulation was obtained by low-pass filtering the CST at 5 Hz. The alpha-and beta-band modulations were obtained by band-pass filtering the CST (8-13 Hz and 13-30 Hz, respectively), extracting the Hilbert envelope, and z-scoring it relative to the normalization window. The synchronisation index was estimated from pairwise spike-train coincidences using a sliding-window approach. Within a 100-ms window advanced in 10-ms steps, synchronisation was computed for all pairs of motor neurons. For each pair, coincident discharges were defined as occurring within ±6 ms across the two spike trains. This window was chosen to reflect short-term synchrony (Datta & Stephens, 1990). The observed count was normalized by the count expected under independent firing, computed from the firing rates. This normalization reduces the dependence of the synchronisation index on firing rate.

From each profile, we extracted two cue-related features: a trend feature and a burst feature. These features were designed to capture slow post ready-cue modulations and transient post-imperative-cue responses, respectively. First, a fourth-degree polynomial was fitted by least square to each trial-averaged profile over the-3 to +1 s window, excluding the 0 to 500 ms post-imperative-cue interval so that the fit reflected the slow trend without being biased by the transient response. The trend feature was defined as the difference between the mean fitted polynomial value in the post-cue window (0 to 1 s) and that in the-3 to-1 s window. The burst feature was defined as the mean signed residual between the observed profile and the fitted polynomial within the 0 to 500 ms post-imperative-cue window. This window was selected because it encompassed the reaction times observed in the Go condition (Fig. 1). Across the four profiles (low-frequency, alpha-band, and beta-band CST modulations, and synchronisation index), this yielded eight cue-related features (2 features × 4 profiles). We additionally extracted summary statistics within three predefined time windows: Baseline (-2 to-1s), Ready (-1 to 0 s), and Post-Cue (0 to 1 s). For each profile and each window, we computed the mean, minimum, and maximum values, yielding 36 features (4 profiles × 3 windows × 3 statistics). Finally, we extracted five timing features from the Post-Cue window: the times of the maximum and minimum of the low-frequency CST modulation, and the times of the maximum of the alpha modulation, beta modulation, and synchronisation index. In total, 49 features were extracted from both simulated and experimental data for each of the three experimental conditions (DF_NoGo, Hand_NoGo, Hand_Go). These features were used for the second inference stage.

### Priors and training simulations

The first inference stage estimated eight parameters describing motor neuron pool properties and baseline synaptic input: the motor neuron size distribution, the amplitudes of the tonic excitatory drive and the three common fluctuating inputs (low-frequency, alpha-band, and beta-band), the amplitude of the independent input, and the variability of common-input and independent-input weights across motor neurons (Table S1). Features for this stage were derived from baseline motor neuron population behaviour in the experimental data, pooled across participants and conditions: the mean, standard deviation and skewness of the distribution of motor neuron firing rates; the same statistics computed for the distribution of inter-spike-interval coefficients of variation; the standard deviations of the low-frequency; alpha-band and beta-band CST modulations in the Baseline window; and the mean synchronisation in the same window. In the second inference stage, the first-stage posterior was used as the prior for the pool and baseline-input parameters, while six cue-related parameters were drawn from uniform priors: a burst amplitude and a trend slope for the net excitatory drive, the alpha-band input, and the beta-band input.

For both inference stages, prior ranges were selected to ensure that simulations generated feature values spanning the range observed experimentally. For the trend in net excitatory drive, an initial run with a symetric prior placed essentially all posterior mass below zero in every condition. The prior was therefore restricted to [−5, 0] µA/s so that training simulations were concentrated in the region supported by the data. Because this restriction makes a negative trend certain a priori, the posterior for this parameter quantifies the magnitude of the decrease rather than providing evidence for its direction. For the first inference stage, we sampled 600 parameter sets from the priors, each repeated 16 times (9,600 simulations in total). For the second inference stage, each parameter set combined pool and baseline-input parameters drawn from the first-stage posterior with cue-related parameters drawn from uniform priors. We sampled 1,200 parameter sets, each repeated 32 times (38,400 simulations in total). For a given parameter set, the 32 repeats had different input time series and served as analogues of individual trials in the experimental data. Repeats were aggregated to generate parameter-set-level feature summaries, which were then used for training the inference network.

Neural density estimators

Simulated parameter sets and their corresponding features were used to train neural density estimators that approximate the posterior distribution of model parameters given observable features. For the first inference stage, we trained a single neural density estimator implemented as a masked autoregressive flow. For the second stage, we trained an ensemble of five such estimators with different random seeds. In both stages, 10% of the simulations were held out for validation and training stopped after 20 epochs without improvement on them. Parameters were rescaled to [0, 1], and features were z-scored before training and inference. Once trained, the neural density estimators were evaluated on the features extracted from the experimental data. For each experimental condition (DF_NoGo, Hand_NoGo, and Hand_Go), they returned posterior distributions over the inferred parameters, representing the range of parameter values compatible with the observed experimental features (Fig. 4C). Reported posteriors were obtained by pooling posterior samples from the five independently trained estimators. Across the ensemble, the estimators produced highly similar posterior estimates.

Validation and posterior predictive checks

We validated both stages of the simulation-based inference pipeline using held-out simulations with known ground-truth parameters. For each held-out simulated observation, 1,000 posterior samples were drawn, and each parameter was summarized by its marginal posterior mode (i.e. peak of the distribution). Parameter recovery accuracy was quantified as the coefficient of determination (R²) between ground-truth values and the posterior-mode estimates, computed separately for each parameter and validation network. Mean R² was 0.81 ± 0.22 in the first stage (baseline inference) and 0.76 ± 0.24 in the second stage (cue-parameter inference).

Calibration was assessed by comparing the nominal probability mass of posterior credible intervals with the empirical frequency at which these intervals contained the ground-truth parameter values. A well-calibrated posterior should show equal nominal and empirical coverage. We quantified calibration as the expected calibration error, defined as the mean absolute difference between nominal and empirical coverage across credible intervals from 0% to 100%. Mean expected calibration error was 6.8 ± 3.9% in the first stage and 7.2 ± 3.4% in the second stage.

Posterior predictive checks were used to assess whether parameters drawn from the inferred posterior could reproduce the experimental data. For each experimental condition, we drew 50 parameter sets from the posterior and simulated 32 trials of motor neuron spiking activity with each set. Figure 4B compares the resulting simulated profiles with experimental condition averages. Each posterior predictive simulation was also reduced to the same 49-element feature vector used for inference and compared with the experimental feature vector. Features were first normalized by the mean absolute value of the corresponding experimental feature across participants. Agreement was summarized as the Pearson correlation coefficient between the posterior-predicted (simulated) feature vector and the experimental feature vector. Mean correlations across posterior predictive simulations were 0.78 ± 0.08 (DF_NoGo), 0.77 ± 0.08 (Hand_NoGo), and 0.75 ± 0.09 (Hand_Go) (Fig. 4B).

## Statistics

Except for simulation-based inference analyses, all statistical analyses were performed in RStudio (Version 2026.01.1+403, Posit Software, Boston, MA, USA). Statistical significance was set at p ≤ 0.05. Data are reported as mean ± standard deviation unless otherwise stated. To quantify task-related modulation of oscillatory activity, alpha-and beta-band power were analysed using linear mixed-effects models implemented with the nlme package. Separate models were fitted for alpha-and beta-band power. Fixed effects included Time (Baseline, Ready, Post-cue), Task (DF_NoGo, Hand_NoGo), and Signal source (EEG vs CST), together with their interaction terms. Participant was included as a random intercept to account for repeated measures and between-participant differences in overall alpha-or beta-band power. To account for heteroscedasticity across time windows, models included a varIdent variance structure (weights = varIdent(form = ∼1 | Time)), allowing a separate residual variance to be estimated for each level of Time. Models with different combinations of interaction terms, including models containing main effects only, were fitted using maximum likelihood estimation. We selected the best-supported models using likelihood-ratio tests and Akaike and Bayesian information criteria, then refitted them using restricted maximum likelihood estimation. When significant main effects or interactions were identified, pairwise post-hoc comparisons were performed using estimated marginal means with Holm correction for multiple comparisons.

Complementary analyses were performed using the same linear mixed-effects modelling framework. First, alpha-and beta-band power derived from CST and conventional bipolar EMG signals were compared during the DF_NoGo and Hand_NoGo conditions. Second, alpha-and beta-band power derived from CST signals were compared between the Hand_NoGo and Hand_Go tasks in the subset of participants for whom EMG decomposition was successful during the Hand_Go condition.

Coherence between the two cumulative spike trains was analysed within the same framework as that described above, with separate models for Fisher-transformed delta-, alpha-and beta-band coherence. Fixed effects were Time, Task and their interaction. Post-hoc comparisons were performed between time windows, with Holm correction.

To complement the window-based analyses and mitigate the limitations associated with temporal averaging, we compared the DF and Hand No-Go conditions using one-dimensional statistical parametric mapping (Pataky, 2010) applied to the continuous time courses of alpha-and beta-band power (n=15). Specifically, a paired SPM{t} was computed over the-2 to 1 s interval, separately for EEG and CST signals. Inference was performed using a two-tailed non-parametric paired test with 5,000 permutations (seed = 1), and cluster-level inference was based on the maximum cluster integral. This analysis was used to identify time intervals during which the temporal evolution of oscillatory power differed between tasks.

## Results

### Task performance

Following an imperative visual cue, participants either performed a ballistic dorsiflexion (DF_Go), a ballistic handgrip (Hand_Go), or no ballistic response (DF_NoGo and Hand_NoGo), while maintaining the baseline submaximal dorsiflexion torque throughout. For Go conditions, participants were instructed to initiate the ballistic response as quickly as possible after the cue. Mean reaction times were 353 ms (95% CI: 329-377 ms) for DF_Go and 370 ms (95% CI: 351-391 ms) for Hand_Go (Fig. 1).

This experimental design allowed us to investigate modulations of alpha and beta oscillatory activity at both the cortical and motor neuron population levels, including in a motor neuron pool not directly involved in the instructed response, namely the TA motor neurons during the handgrip task.

Identification of motor neuron spiking activity

Identifying motor neuron spiking activity was necessary because conventional bipolar EMG lacked sufficient sensitivity to reliably detect alpha-and beta-band modulations, as shown in complementary analyses (Fig. S1).

Decomposing high-density surface EMG during ballistic contractions remains technically challenging. Consequently, TA EMG signals from the DF_Go task (in which the TA itself produced the ballistic response) could not be reliably decomposed and were not included in the analyses. Consistent with our primary aim of testing whether experimentally induced alpha-and beta-band modulations are expressed in spinal motor neuron output, motor neuron analyses focused on the No-Go conditions, where such modulations have been reported to be particularly prominent (Zicher *et al*., 2024). For the Hand_Go task, in which the TA was not directly involved in the ballistic response, EMG signals were successfully decomposed in a subset of participants (n = 10/15) who maintained dorsiflexion at the required submaximal torque for at least 200 ms after the ballistic handgrip. In the remaining participants, dorsiflexion torque was released around the time of the handgrip response, leaving insufficient data for reliable time-frequency analyses over a meaningful time window. Motor unit data from the Hand_Go task are therefore presented as complementary analyses to inform the interpretation of the results and support the discussion.

Across the three analysed tasks (DF_NoGo [n=15], Hand_NoGo [n=15], and Hand_Go [n=10]), a total of 5,963 motor neuron spike trains were identified. Each spike train corresponded to a single motor neuron tracked across all 13 trials of a given condition (No-Go or Go) within a run. The mean number of motor neurons identified per run and per participant was 36.0 ± 9.5 for DF_NoGo (range: 17-55), 36.9 ± 9.0 for Hand_NoGo (range: 15-53), and 39.6 ± 10.2 for Hand_Go (range: 22-57). Of note, decomposition was performed independently for each run and motor units were not tracked across runs. A motor unit identified in several runs of the same task was therefore counted once per run, so the numbers reported above reflect motor neurons identified per run rather than unique motor neurons per participant.

Alpha-and Beta-band modulations

To assess modulations of power in the alpha (8-12 Hz) and beta (13-30 Hz) bands, we computed time-frequency maps from both EEG signals and the summed spiking activity of identified TA motor neurons, referred to as cumulative spike train (CST). For EEG, time-frequency maps were averaged over a cluster of electrodes (Cz, C1, C3) covering the expected medial and lateral sensorimotor regions corresponding to leg and hand representations. The scalp distribution showed alpha-and beta-band activity over this cluster during No-Go conditions, together with beta activity over frontal areas (Fig. 2), consistent with previous reports using methods with higher spatial resolution (Schaum *et al*., 2021).

**Figure 2.**
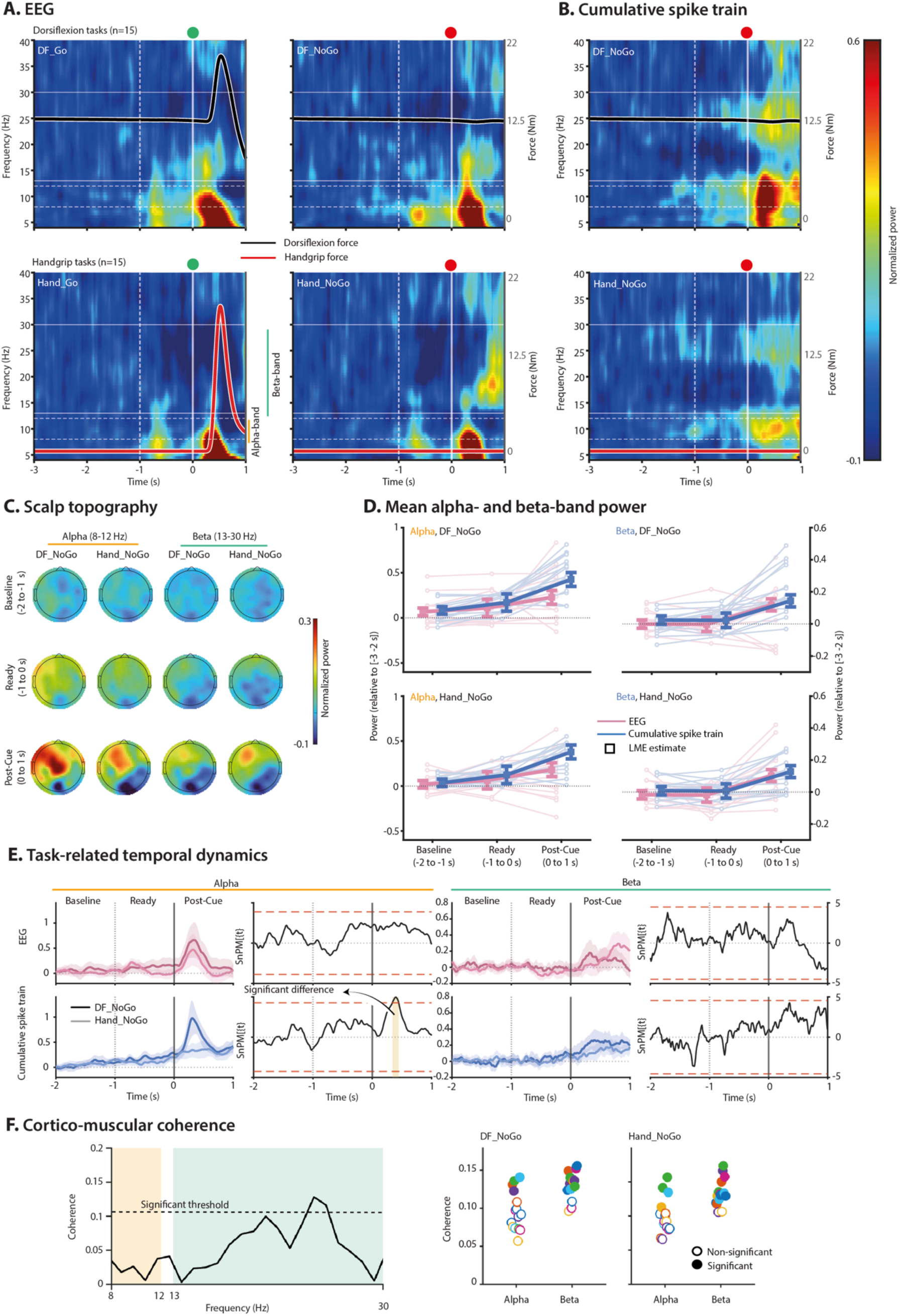
Alpha-and beta-band oscillatory dynamics in electroencephalography (EEG) signals and motor neuron output. (A, **B)** Time–frequency maps of normalized power, computed across tasks for EEG signals (**A**, averaged over Cz, C1, and C3) and cumulative spike trains (**B**, CST) across tasks. The dashed vertical line indicates the Ready cue, and the solid vertical line indicates the imperative cue (Go or No-Go). Torque traces are overlaid. **(C)** Scalp topographies of alpha (8–12 Hz) and beta (13–30 Hz) power across the Baseline (-2 to-1), Ready (-1 to 0), and Post-Cue (0 to 1 s) windows. **(D)** Individual mean alpha-and beta-band power values across time windows, with the linear mixed-model estimates (± standard error) overlaid as filled markers. Alpha power increased from Baseline to Ready and Post-Cue in CST, whereas EEG alpha increased only Post-Cue, with overall higher alpha power in DF_NoGo than Hand_NoGo. Beta power increased Post-Cue in both EEG and CST signals, with no Signal source-dependent effect. **(E)** Time-varying alpha-(left) and beta-band (right) modulations for EEG (top) and CST (bottom). Shaded area indicates the 95% confidence interval across participants. Statistical parametric mapping revealed a larger post-cue alpha peak in CST signals for DF_NoGo than Hand_NoGo, whereas beta-band activity showed similar dynamics across tasks. **(F)**. Representative coherence function for an individual participant (left panel) with maximal coherence values represented for each participant in the right panels. Data are presented for 15 participants. DF, dorsiflexion.

We first estimated normalized power within 1-s windows, selected to match previous work (Zicher *et al*., 2024) and to balance temporal precision, spectral reliability, and signal-to-noise ratio for estimating alpha-and beta-band power. The analysis focused on three windows: Baseline (−2 to −1 s), Ready (−1 to 0 s), and Post-Cue (0 to 1 s). In the alpha band, the linear mixed-effects model revealed a significant Time × Signal source interaction (F(2, 159) = 3.63, p = 0.029; Fig. 2). For CST signals, normalized alpha power showed clear temporal modulation (Fig. 2), with significant increases at both Ready (-1s to 0 s) and Post-Cue (0 s to 1 s) compared to Baseline (-2 s to-1 s; all p < 0.002), as well as a significant increase from Ready to Post-Cue (p < 0.001). EEG-derived alpha power followed a similar pattern but increased only from Baseline and Ready to Post-Cue (p = 0.003 and p = 0.03, respectively), with no significant difference between Baseline and Ready (p = 0.12). A main effect of Task was also observed (p = 0.013), with overall higher alpha power during DF_NoGo than Hand_NoGo.

In the beta band, model comparison indicated that interaction terms, including those involving Signal source, did not improve the model fit. The final model revealed a main effect of Time (F(2, 161) = 19.08, p < 0.001), indicating similar temporal modulation across EEG and CST signals. Specifically, beta power was significantly higher at Post-Cue than at both Baseline and Ready (both p < 0.001; Fig. 2), with no difference between Baseline and Ready (p = 0.89). These results indicate that beta power increased after the cue in both cortical activity and TA motor neuron output.

To complement the window-based analyses and improve temporal resolution, we applied one-dimensional statistical parametric mapping to the continuous time-course of alpha-and beta-band power, focusing on differences between tasks (DF_NoGo vs. Hand_NoGo). In CST signals, alpha power was higher in DF_NoGo than in Hand_NoGo over a single cluster spanning 0.34 to 0.45 s after the imperative cue (t* = 4.19; p = 0.002). No between-task differences were found for alpha power in EEG signals (t* = 3.83), or for beta power in either CST (t* = 4.54) or EEG signals (t* = 4.42). These results suggest that alpha-band modulation is larger in the responding motor neuron pool, consistent with a more focal expression, whereas beta-band modulation showed no such task dependence, consistent with a more widespread expression.

In the subset of participants with available motor neuron data in the Hand_Go task (n=10), we performed complementary analyses comparing alpha-and beta-band CST modulations between Hand_NoGo and Hand_Go (Fig. S2). For both frequency bands, model comparison supported an additive model including Condition (Go and No-Go) and Time without interaction. Alpha power did not differ between conditions (F(1,47) = 0.15; p = 0.70) but varied across time-windows (main effect of Time: F(2,47) = 14.33; p < 0.001), with higher values at Post-Cue than at Baseline (p < 0.001) and Ready (p = 0.035); and higher values at Ready than at Baseline (p = 0.035). Beta power was higher in No-Go than Go trials (main effect of Condition: F(1,47) = 4.02; p = 0.050) and varied across time windows (main effect of Time: F(2,47) = 9.10, p < 0.001), with higher values at Post-Cue than at both Baseline and Ready (both p < 0.001), and no difference between Baseline and Ready (p = 0.72). These exploratory analyses suggest that alpha-and beta-band modulations were also present in TA motor neuron output during handgrip execution, when TA remained tonically active but was not involved in the motor response. Together, these results provide direct evidence that task-related alpha-and beta-band modulations are expressed in a motor neuron pool beyond those involved in response selection, suggesting that cortical oscillatory dynamics extend beyond the responding effector.

Oscillatory activity shared across motor neurons

To determine whether the alpha-and beta-band modulations observed during the No-Go trials reflected a change in the oscillatory activity shared across motor units rather than the accompanying change in discharge rate, we computed the coherence between the cumulative spike trains of two disjoint subsets of motor units, in the same three windows as for time-frequency maps (Fig. 3). Delta-band coherence did not differ between tasks and showed no Time × Task interaction, and the effect of Time did not reach significance (Task: F(1, 72) = 0.00, p = 0.996; Time: F(2, 72) = 2.89, p = 0.062). In the alpha-band, the linear mixed-effects model revealed a significant Time × Task interaction (F(2, 70) = 3.59, p = 0.033). In DF_NoGo, alpha-band coherence increased from Baseline and Ready to Post-Cue (both p < 0.001), with no difference between Baseline and Ready (p = 0.23). In Hand_NoGo, alpha-band coherence did not change significantly across time-windows (all p > 0.087). At Post-Cue, alpha-band coherence was higher in DF_NoGo than in Hand_NoGo (p = 0.020), whereas it did not differ between tasks at Baseline or Ready (both p = 0.68). In the beta band, the model also revealed a significant Time × Task interaction (F(2, 70) = 3.56, p = 0.034). Beta-band coherence increased from Baseline and Ready to Post-Cue in both DF_NoGo (both p < 0.001) and Hand_NoGo (both p = 0.003), with no difference between Baseline and Ready (p = 0.056 and p = 0.92, respectively). Although absolute coherence at Post-Cue did not differ significantly between tasks (p = 0.070), the increase from Baseline to Post-Cue was larger in DF_NoGo than in Hand_NoGo (+44% vs +20%; t(14) = 5.04, p = 0.0004). Over the same time-windows, discharge rate decreased slightly from Baseline to Post-Cue, by a similar amount between tasks (DF_NoGo: −0.55 ± 0.41 pps; Hand_NoGo: −0.58 ± 0.44 pps; t(14) = 0.28, p = 0.78).

**Figure 3.**
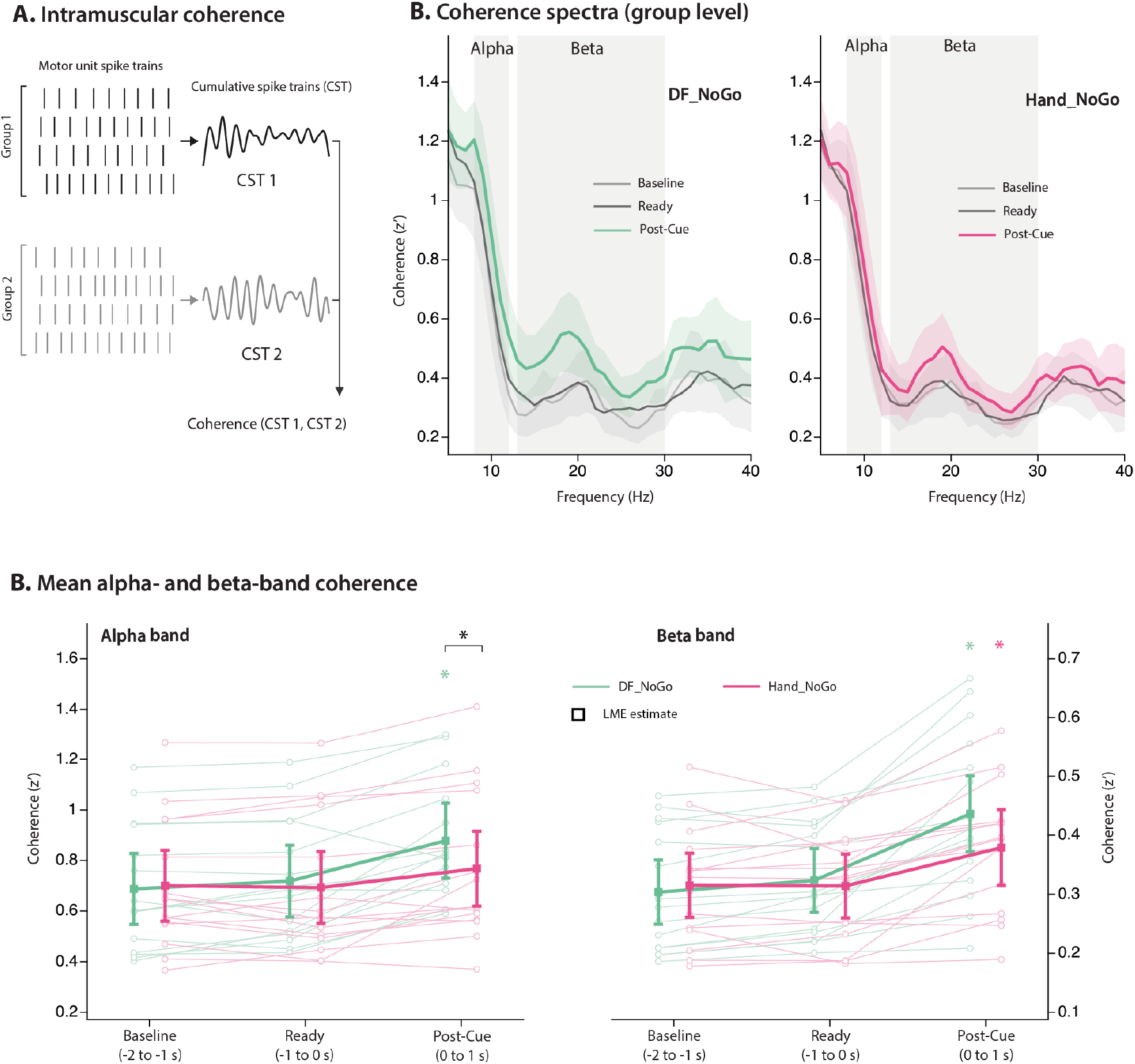
Coherence between the spike trains of two disjoint motor unit subsets. **(A)** For each run, the identified motor units were split at random into two disjoint subsets of equal size (k = half of the motor neurons identified in that run), each subset was summed into a cumulative spike train (CST), and coherence was computed between the two resulting CSTs. **(B)** Coherence spectra (Fisher-transformed, z′) for DF_NoGo (left) and Hand_NoGo (right), in the three time-windows, with the shaded areas indicating the 95% confidence interval across participants. **(C)** Individual alpha-and beta-band coherence values (n=15) across time windows (thin lines), with the linear mixed-model estimates (± 95% confidence interval) overlaid as filled markers. Alpha-band coherence increased from Baseline and Ready to Post-Cue in DF_NoGo only. Beta-band coherence increased from Baseline and Ready to Post-Cue in both tasks. Alpha-band coherence was higher at Post-Cue for DF_NoGo than Hand_NoGo. CST, cumulative spike train.

**Figure 4.**
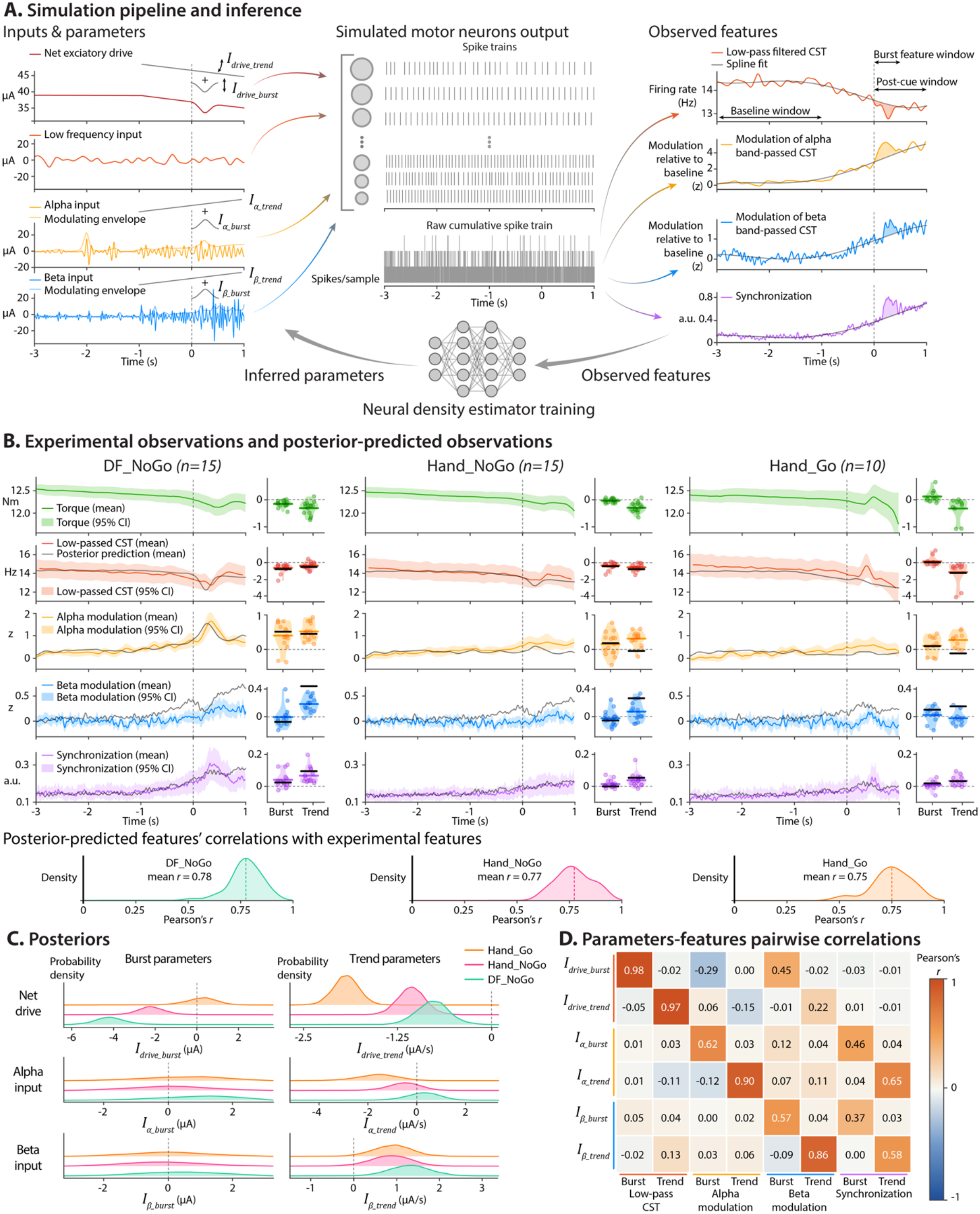
Simulation-based inference of synaptic inputs to motor neurons. **(A)** Simulations of motor neuron spiking activity were driven by four input components: a net excitatory drive, together with three zero-mean band-limited Gaussian fluctuating inputs in the low-frequency (< 5 Hz), alpha, and beta ranges. Cue-related modulations of the net excitatory drive, alpha-band and beta-band inputs were modelled as a transient burst and a linear trend (see Methods). The resulting spike trains were summed to obtain the cumulative spike train (CST), from which four output traces were computed: the low-pass filtered CST, alpha-and beta-band CST modulations, and motor neuron synchronisation. Observable features were extracted to quantify cue-related modulations. Note that eight representative features are shown, while 49 features in total were used for inference. A neural density estimator was trained to map these features to the burst and trend input parameters. **(B)** Experimental observations (colored traces; mean ± 95% CI across participants) and posterior-predictive simulations (black; mean across simulations) are depicted for each condition. Burst and trend feature distributions are shown to the right, with each dot representing a participant; experimental and simulated means are indicated by colored and black lines, respectively. Bottom: distributions of Pearson correlations between predicted and observed features across 50 posterior-predictive simulations per condition (see Methods). **(C)** Posterior distributions of the six inferred input parameters (burst and trend parameters for net excitatory drive, and alpha-and beta-inputs) for each condition. **(D)** Pairwise correlations between input parameters (y-axis) and observable features (x-axis; 8 out of the 49 features) across the training simulation set. DF, dorsiflexion.

In the subset of participants with Hand_Go data (n = 10), delta-and alpha-band coherence did not differ between conditions (Hand_Go vs. Hand_NoGo) or across windows. In contrast, model comparison retained an effect of Time for beta-band coherence (F(2, 48) = 12.50, p < 0.001), with no effect of Condition and no interaction. Averaged across conditions, coherence was higher at Post-Cue than at both Baseline and Ready (both p < 0.001), with no difference between Baseline and Ready (p = 0.58).

Together, these results show that beta oscillations shared across TA motor neurons increased after the imperative cue in both tasks, including when the muscle was not involved in the response, although the increase was larger when it was. In the alpha band, the increase in coherence was significant only when the TA was the responding muscle.

Effects on force, low-frequency drive and motor neuron synchronisation

To determine whether task-related oscillatory modulations were accompanied by changes in motor output, we analysed dorsiflexion torque, the low-frequency component of CST (5 Hz cut-off), and motor neuron synchronisation across all available data (DF_NoGo, n = 15; Hand_NoGo, n =15; Hand_Go, n = 10). For each signal, we extracted two features: a burst feature, defined as the mean value within a post-cue window (0–500 ms for CST and synchronisation; 200–700 ms for torque to account for electromechanical delay (Del Vecchio *et al*., 2018)), and a trend feature, defined as the post-cue mean (0 to 1 s) minus the reference period (-3-1 s; see Methods). Each feature was tested against zero using bootstrap 95% confidence intervals (CI). Features were considered significant when their CI excluded zero. Torque showed a significant negative burst only in DF_NoGo (CI [−0.14 to −0.07] Nm, Fig. 4B). Low-frequency CST bursts were negative in DF_NoGo (CI [−0.96 to −0.44] pps) and Hand_NoGo (CI [−0.58 to −0.18]) but not in Hand_Go (CI [−0.17 to 0.56]). Synchronisation bursts were positive in DF_NoGo (CI [0.019 to 0.069]) and Hand_NoGo (CI [0.003 to 0.027]), with no clear burst in Hand_Go (CI [−0.010 to 0.026]).

Trends were more consistent across tasks. Torque and CST trends were negative in all three conditions, whereas synchronisation trends were positive. Specifically, torque decreased in DF_NoGo (CI [−0.28 to −0.13]), Hand_NoGo (CI [−0.24 to −0.13]), and Hand_Go (CI [−0.38 to −0.08]). CST showed the same pattern in DF_NoGo (CI [−0.597 to −0.192]), Hand_NoGo (CI [−0.799 to −0.397]), and Hand_Go (CI [−2.227 to −0.428]). Conversely, synchronisation increased in DF_NoGo (CI [0.046 to 0.096]), Hand_NoGo (CI [0.021 to 0.062]), and Hand_Go (CI [0.016 to 0.057]).

Together, these results indicate a systematic modulation of motor output in DF_NoGo, characterized by reduced torque and firing rate (low-pass filtered CST) together with increased motor neuron synchronisation. The handgrip tasks showed less systematic but still significant modulations, especially for the trend features. These effects should nevertheless be interpreted cautiously for Hand_Go given the smaller sample size (n = 10).

Coherence between EEG and CST

To assess frequency-specific coupling between cortical and motor neuron oscillations, we computed coherence between each EEG channel in the predefined sensorimotor cluster (Cz, C1, C3) and the CST over the Post-Cue window. Coherence was considered detectable when at least one frequency bin within the band of interest exceeded the upper 95% confidence limit. This analysis was used to determine whether alpha-and beta-band modulations in the CST were accompanied by measurable coupling with cortical activity.

In the beta band, detectable EEG-CST coherence was observed in 13 of 15 participants during Hand_NoGo and 12 of 15 during DF_NoGo, although coherence values were relatively low (Fig. 2F), consistent with typical cortico-muscular coherence estimates from non-invasive recordings (Negro & Farina, 2011; Abbagnano *et al*., 2025). In the alpha band, coherence was detected in only 4 of 15 participants, consistent with previous work showing that alpha-band coherence is relatively rare (Baker, 2007). Several factors may explain this difference. First, alpha modulations in our data appeared to be more transient than beta modulations (Fig. 2), which may attenuate coherence estimates computed over an extended 1-s window. Second, even if cortical alpha oscillations are reliably transmitted to motor neurons, spinal-level processes such as recurrent inhibition (Williams & Baker, 2009) may interfere with this propagation, weakening cortico-spinal coherence. Third, the low prevalence of alpha-band coherence may reflect that part of the alpha-band modulation observed in motor neuron output originates from non-cortical sources, including subcortical or brainstem pathways (Grosse *et al*., 2003).

### Simulation-based inference

Experimental data revealed task-related changes in motor output (torque, low-frequency CST, synchronisation, and alpha-and beta-band power), but these observations alone cannot identify which synaptic input components account for them. Multiple input components, such as net excitatory drive and alpha-and beta-band inputs, may contribute simultaneously and interact in such a way that different combinations of inputs can produce similar observable changes in motor output. Here, we used simulation-based inference, which estimates posterior distributions over model parameters (neural inputs in our case) by training a neural network on simulations from a generative model of motor neuron output. This approach is particularly useful for models of spiking neurons, where it is difficult or impossible to directly calculate how likely a given input is to generate the observed activity. Instead of identifying a single ‘best’ solution, simulation-based inference estimates a range of plausible input configurations and indicates how likely each one is. This makes it possible to quantify uncertainty and identify the inputs that most likely explain the experimental observations. Of note, our pipeline was validated on simulated data with known ground truth (Fig. 4B), as recommended (Tolley *et al*., 2024) (see Methods).

When considering the modulation of net excitatory input, a clear pattern emerged. In the No-Go conditions, posterior distributions for the post-cue burst parameter lay entirely below zero (Fig. 4C), providing strong evidence for a transient decrease in excitatory drive after the imperative cue. This decrease was more pronounced in DF_NoGo than in Hand_NoGo (pΔ = 0.97; i.e., 97% posterior probability that DF_NoGo > Hand_NoGo). In addition to these transient modulations, net excitatory drive decreased progressively over the −1 to 1 s window in all tasks. Of note, because the prior for this parameter was restricted to non-positive values (see Methods), the posterior quantifies the magnitude rather than the direction of change.

The alpha-and beta-band inputs showed distinct patterns. For beta-band input, the post-cue burst parameter was centred near zero in all conditions (Fig. 4C), whereas the trend parameter showed a high probability of an increase in all three tasks (all P(θ > 0) > 0.96), indicating an increase in beta-band input regardless of whether the TA was involved in the response. For alpha-band input, the only evidence for an increase was in DF_NoGo, with a moderate probability for the burst (P(θ > 0) = 0.77) and a weak one for the trend (P(θ > 0) = 0.67). In the handgrip tasks, by contrast, the alpha-band trend was more likely negative (Hand_Go: P(θ < 0) = 0.98; Hand_NoGo: P(θ < 0) = 0.78), with burst distributions centred near zero. Together with the modulation of net excitatory drive described above, these results indicate that the Go/No-Go task engaged band-specific changes in oscillatory inputs.

To further dissociate the contributions of individual input components, we examined the marginal correlations between simulation parameters and the resulting feature values across the training set (Fig. 4D). Parameter sets were sampled from the prior independently from one another, so parameter values were uncorrelated across the training set. As such, these marginal correlations are not confounded by covariance between parameters and instead reflect the sensitivity of each feature to variation in the corresponding input. The low-pass filtered CST burst feature was strongly and positively associated with the transient burst of net excitatory drive (r=0.98), but not with alpha-or beta-band inputs (all |r| ≤ 0.05). This supports the interpretation that the post-cue decreases in population firing rate (low-pass CST) observed in the No-Go conditions primarily reflected modulations in the net excitatory drive to motor neurons. In contrast, net excitatory drive showed no relationship with motor neuron synchronisation (|r| ≤ 0.03). Instead, post-cue synchronisation was more strongly associated with alpha-and beta-band inputs, including both transient burst parameters (r=0.46 for alpha; r=0.37 for beta) and temporal trend parameters (r=0.65 for alpha; r=0.58 for beta). This dissociation suggests that net excitatory drive predominantly shapes the overall level of motor output, whereas higher-frequency inputs shape the temporal synchronisation of motor neuron discharges.

## Discussion

Using large-scale recordings of spinal motor neuron populations, this study provides evidence that beta-band oscillations during a Go/No-Go task are expressed in a motor neuron pool beyond that directly involved in the instructed response. This suggests a broader expression of higher-frequency oscillations within the peripheral motor system than is typically assumed. Our simulations suggest that these oscillatory inputs account for a transient increase in motor neuron synchronisation but do not directly explain the significant, albeit weak, changes in motor output (torque). Instead, simulation-based inference indicates that changes in motor output are primarily explained by modulations in net excitatory drive, possibly reflecting changes in corticospinal excitability. Overall, these findings may be consistent with a parallel control architecture in which low-frequency drive controls motor output, while higher-frequency inputs broadly shape motor neuron synchronisation. One speculative possibility is that such inputs help poise the whole motor system for rapid adjustments in output.

The Go/No-Go paradigm was chosen because it reliably induces cortical alpha-and beta-band modulations, with successful inhibition marked by a post-cue power increase (Pani *et al*., 2014; Zicher *et al*., 2024). Consistent with this, both tasks produced the expected alpha-and beta-band power increases after the imperative cue (Fig. 2), confirming that the paradigm effectively modulated the cortical oscillatory dynamics that we aimed to track in spinal motor neuron output. Because oscillatory activity in peripheral neural output can only be assessed when motor neurons are active, we focused on the TA muscle during submaximal contractions, in tasks where it was either the responding (DF_Go/NoGo) or a non-responding muscle (Hand_Go/NoGo). Analyses were primarily restricted to the No-Go conditions because motor unit decomposition is not sufficiently reliable during ballistic contractions. Importantly, this motor neuron-based approach enabled detection of subtle oscillatory modulations that are obscured in conventional interference EMG signals (Fig. S1).

We observed an increase in alpha-and beta-band power in TA motor neurons during both tasks, including the Hand_NoGo, despite a smaller post-imperative-cue alpha peak than during DF_NoGo (Fig. 2E). The beta-band modulation in the non-responding muscle is partly in line with a recent observation made during movement cancellation, in which beta power in the non-responding muscle increased with a moderate effect size without reaching significance, a result interpreted as effector-specific transmission (McGeady *et al*., 2026). One explanation for the absence of a significant change in that study is that beta power was derived from interference EMG, which is less sensitive than motor neuron output to reveal beta-band oscillations (Fig. S1). The discrepancy between our study and that of McGeady et al. (2026) may therefore reflect a difference in measurement sensitivity rather than a genuine difference in the underlying phenomenon. Regardless of its statistical significance, however, the modulation reported in McGeady et al. (2026) is difficult to interpret in terms of how broadly beta-band input is distributed across the motor system. In that study, the non-responding muscle was the homologous wrist extensor of the contralateral arm, and both arms maintained the contraction. Because homologous muscles of the two limbs can receive shared descending input (Carson, 2005), a beta modulation in the non-responding arm could arise from this shared input. A muscle that is neither the homologue of the responding muscle nor located in the same limb therefore provides more direct evidence of broad distribution within the motor system. Here, beta-band power in TA motor neurons increased while the response was produced by the hand, indicating that this modulation is neither restricted to the responding muscle nor to its contralateral homologue.

Interpreting modulations of CST band power as changes in oscillatory input requires an important consideration. The CST power spectrum contains a strong component at the mean discharge rate, so the post-cue decrease in discharge rate could alone increase power in the alpha band. For this reason, we used coherence between disjoint subsets of motor units to assess the oscillatory activity that is shared across motor neurons, as this measure is much less sensitive to changes in discharge rate. Coherence changed around the imperative cue in a band-dependent manner. Specifically, beta-band coherence increased whether or not the TA was involved in the response, although the increase was larger when it was. This suggests that the shared beta oscillations were not restricted to the responding motor pool, while remaining weighted towards it. In contrast, alpha-band coherence increased only when the TA was the responding muscle, indicating a more selective distribution. To our knowledge, this is the first report of a cue-related modulation of alpha-band oscillatory activity in the responding muscle, since previous work on movement cancellation focused on the beta and low-gamma bands (Zicher *et al*., 2024). Whether these shared oscillations can be explained by changes in common synaptic input was then tested with simulation-based inference (Fig. 4). Specifically, inference analysis revealed a high probability of a positive trend in beta-band input across all three conditions (DF_NoGo, Hand_NoGo, and Hand_Go), i.e., regardless of whether TA was involved in the response. Such a pattern is consistent with the view that beta-band activity reflects a nonselective adjustment of motor output and may correspond to a global “pause-like” process (Wessel, 2020). Importantly, in the *pause-then-cancel* model, the Pause is not specific to stopping and can also be engaged after a Go cue (Diesburg & Wessel, 2021), consistent with the presence of a post-cue beta increase in Go trials (Fig. S2). The alpha-band input behaved differently with the only evidence for an increase being the post-cue burst in DF_NoGo, mirroring the coherence results.

Our results support the conclusion that cue-related oscillatory modulations reflect changes in the common input to TA motor neurons, in both tasks for beta and only in DF_NoGo for alpha. The origin of these modulations, however, remains an open question. Even though it is probable that alpha-and beta-band modulations in spinal motor neuron output reflect transmission of cortical signals, an alternative possibility is that they originate from non-cortical mechanisms such as neuromodulation of brainstem/spinal neurons (Thompson *et al*., 2019) or represent parallel manifestations of a common process acting at both cortical and spinal levels, rather than direct top-down transmission. For example, beta-band oscillations can be generated by spinal circuits such as recurrent inhibition (Dernoncourt *et al*., 2026). Furthermore, alpha-band common input is classically attributed to oscillations within the spindle feedback loop (Christakos *et al*., 2006). The selective increase we observed could therefore reflect a change in fusimotor drive or reflex gain in the muscle prepared for the action rather than a descending oscillatory input. Nevertheless, several converging lines of evidence support the transmission hypothesis. First, the Go/No-Go paradigm used here is known to engage specific cortical alpha-and beta-band oscillatory dynamics, and we observed highly similar modulations at the cortical (EEG) and spinal (CST) levels. Second, significant beta-band corticomuscular coherence (albeit weak) was observed in most participants (up to 13/15). Third, a recent study supports the existence of a functional corticospinal pathway, showing that evoking cortical beta oscillations with subthreshold transcranial magnetic stimulation elicits a robust, short-latency, time-locked response in spinal motor neurons (Sarasquete *et al*., 2026), and rhythmic subthreshold stimulation of the motor cortex is transmitted approximately linearly to the motor neuron pool, most effectively in the 20–50 Hz range (Cabral *et al*., 2026). Together, these observations support the interpretation that the oscillatory activity in motor neuron output reflects the transmission of cortical signals to the periphery. We nonetheless acknowledge that coherence values were modest, as commonly reported, and that further work is needed to provide direct evidence for brain-to-muscle transmission (or muscle-to-brain transmission).

The presence of alpha and beta common input at the level of motor neuron populations raises a fundamental question about the functional significance of transmitting signals that may not directly influence mechanical output. The prevailing view is that beta-band oscillations (Farina *et al*., 2016; Zicher *et al*., 2023; Ibanez *et al*., 2025), and to a lesser extent alpha-band oscillations (Zicher *et al*., 2023), lie within a “motor null space” with respect to force production. This view is grounded in the mechanical properties of the musculoskeletal system, often modelled as a second-order low-pass filter that strongly attenuates the mechanical effects of higher-frequency neural inputs, typically above ∼10 Hz (Ibanez *et al*., 2025). However, whereas previous studies generally considered these oscillations as tonic inputs, emerging evidence suggests that burst-like beta-band inputs may have greater functional impact by transiently synchronizing subpopulations of motor neurons (Pascual Valdunciel *et al*., 2026), and, under some conditions, inducing small force fluctuations when their amplitudes reach physiological extremes (Watanabe & Kohn, 2015; Zicher *et al*., 2023). The Go/No-Go paradigm used here was designed to elicit such burst-like activity, allowing us to test whether alpha-and beta-band modulations are expressed in spinal motor neuron output under conditions in which these signals may be of larger amplitude than those typically observed during sustained contractions. As shown in Fig. 4B, participants exhibited systematic, albeit small, changes in population firing rate (low-pass CST) and torque, comprising both transient post-cue bursts and slower trends, with effects more pronounced during DF_NoGo. However, our simulation-based inference indicates that alpha-and beta-band inputs do not account for these changes. Instead, these changes were best explained by modulations in net excitatory drive. Specifically, posterior distributions supported a decrease in excitatory drive after the No-Go cue (with a larger decrease when the TA muscle was the responding muscle) and an increase in excitatory drive during Hand_Go (Fig. 4C). This pattern aligns with previous work showing that response preparation is accompanied by broad, non-selective changes in corticospinal excitability that can extend to task-irrelevant muscles, while often being more pronounced in task-relevant muscles (Greenhouse *et al*., 2015). Such broad modulations may be computationally less demanding than selective, effector-specific targeting, providing a robust default state that can be rapidly refined once the required action is specified and flexibly adapted during ongoing movement.

Although alpha-and beta-band inputs did not primarily explain changes in low-frequency drive, our simulations highlight their role in shaping motor neuron synchronisation (Fig. 4D). This aligns particularly well with previous simulation work showing that short-term synchronisation increases when the input frequency approaches the motor units’ own mean discharge rate, a range that, in many muscles, overlaps with the alpha and low-beta bands (Lowery & Erim, 2005). Whether this effect on synchronisation serves any functional purpose, however, remains unclear. It could simply be an unavoidable by-product of alpha-and beta-band oscillations reaching the motor neuron pool, without any adaptive role, or it could reflect a mechanism by which the nervous system shapes the functional state of the motor neurons. Consistent with this latter possibility, precise spike timing, beyond discharge rate alone, has been shown to carry functionally important information for motor output across species and levels of the motor system (Sober *et al*., 2018). Motor neuron synchronisation has been suggested to facilitate rapid force production (Semmler, 2002), but direct evidence remains limited. Of note, motor neuron synchronisation was also observed during Hand conditions (when the TA was not the response effector), indicating that this effect is broadly expressed across active motor pools, although its amplitude was larger when the TA was the responding muscle. Together, these findings provide a plausible mechanism by which higher-frequency oscillatory inputs could prepare the motor system for rapid changes in motor output, although this remains to be tested experimentally. They also refine the notion of a motor null space: although alpha-and beta-band oscillations do not directly control mean force, they are not functionally silent. By shaping motor neuron synchronisation, they may indirectly influence the rate at which force changes in response to a given change in net excitatory drive, altering how the motor pool transduces that input rather than contributing to the drive itself.

Why synchronisation-shaping signals might be broadly transmitted rather than restricted to the responding motor neuron pools remains speculative. One possibility is that synchronisation has limited behavioural consequences in motor pools whose net excitatory drive remains unchanged, becoming functionally relevant primarily when drive changes. Broadly increasing synchronisation may therefore impose little cost in non-responding motor pools, reducing the need for precisely targeted modulation. A similarly broad, nonselective mode of preparatory modulation has been described in other contexts, where motor-evoked potentials are suppressed in task-irrelevant, non-homologous muscles alongside a separate effector-specific mechanism, consistent with a general gain-setting process operating independently of more targeted control (Greenhouse *et al*., 2015).

In conclusion, our results show that the alpha-and beta-band oscillatory dynamics elicited by a Go/No-Go task are expressed in spinal motor neuron output more broadly than classically assumed. Together, motor unit coherence and simulation-based inference indicate that these modulations reflect common synaptic input rather than discharge rate. Beta-band common input increased whether or not the muscle was involved in the response, whereas the alpha-band increase was confined to the responding muscle. Rather than altering force, these inputs primarily shaped motor neuron synchronisation. They may reflect a parallel control architecture in which low-frequency modulations of net excitatory drive determine motor output, while higher-frequency inputs set the state of motor pools, though more selectively in the alpha band. The central nervous system may thus multiplex motor commands with state-shaping signals to configure the peripheral motor system for upcoming actions. Future experiments should test this hypothesis against the simpler alternative that these oscillations are transmitted to motor neurons without serving any particular purpose, as a by-product of their generation upstream.

## Supplemental data availability

Code used for modelling and simulation-based inference is available at: https://github.com/FrancoisDernoncourt/Oscillatory_inputs_to_motor_neurons

The full dataset (raw EEG and EMG data and edited motor unit spike trains) will be made publicly available upon publication. During peer review, the data can be accessed via the following temporary link: https://figshare.com/s/25f5c3d7bddbb32c9252

## Funding resources

This study was funded by a grant from the French National Research Agency (ANR-24-ASTR-0003; NeuroCom project). François Hug is supported by the French government, through the UCAJEDI Investments in the Future project managed by the ANR with the reference number ANR-15-IDEX-01, by a fellowship from the *Institut Universitaire de France* (IUF), and by an equipment grant from the *Région Sud*. Wolbert van den Hoorn is supported by a grant from The University of Queensland (new staff research start fund).

## Author Contributions

Contribution and design of the experiment: FH, FD, WvdH; Collection of data: FH, WvdH; Analysis and interpretation: FH, CN, FD, WvdH; Drafting the article or revising it for important intellectual content: FH, CN, FD, WvdH; All authors approved the final version of the manuscript.

---

**Supplementary Figure 1.**
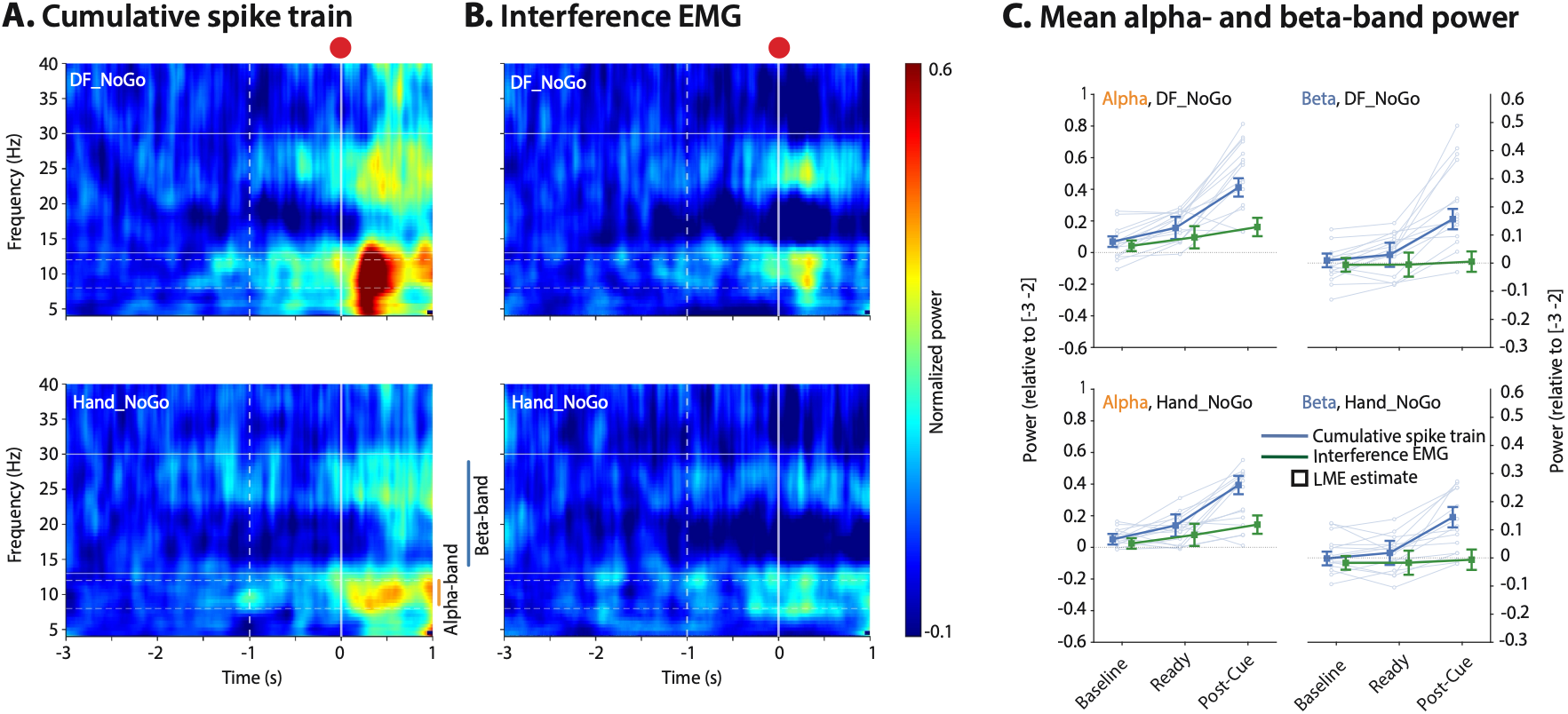
Comparison of cumulative spike train and conventional bipolar electromyography. We assessed the sensitivity of interference electromyography (EMG) signals for detecting modulations in the alpha and beta bands. To this end, we reproduced a classical bipolar electrode configuration using the high-density EMG grid positioned closer to the location typically used for conventional bipolar EMG recordings (for more details, see (Avrillon *et al*., 2021)) and calculated time-frequency maps on unrectified EMG. Linear mixed-effects models revealed a significant Time × Signal source interaction for both alpha (F(2, 159) = 10.21, p < 0.001) and beta bands (F(2, 159) = 11.59, p < 0.001). In the alpha band, cumulative spike train (CST) signals showed clear temporal modulation across all time windows (all p < 0.001), whereas bipolar EMG signals exhibited significant differences only between Baseline and Post-Cue (p = 0.003) and between Baseline and Ready (p = 0.042); Ready and Post-cue showing no significant difference (p = 0.075). In the beta band, CST signals also showed clear temporal modulation, with increased power in the Post-Cue window compared to both Baseline and Ready (both p < 0.001), whereas the EMG signals showed no significant changes (all p = 1). In addition, normalized power in both alpha and beta bands was higher for CST signals than for bipolar EMG signals during the Ready and Post-Cue windows (all p < 0.017). Together, these results indicate that CST provide greater sensitivity than interference EMG for detecting temporal modulations in higher frequency activity, particularly in the beta-band. DF, dorsiflexion.

**Supplementary Figure 2.**
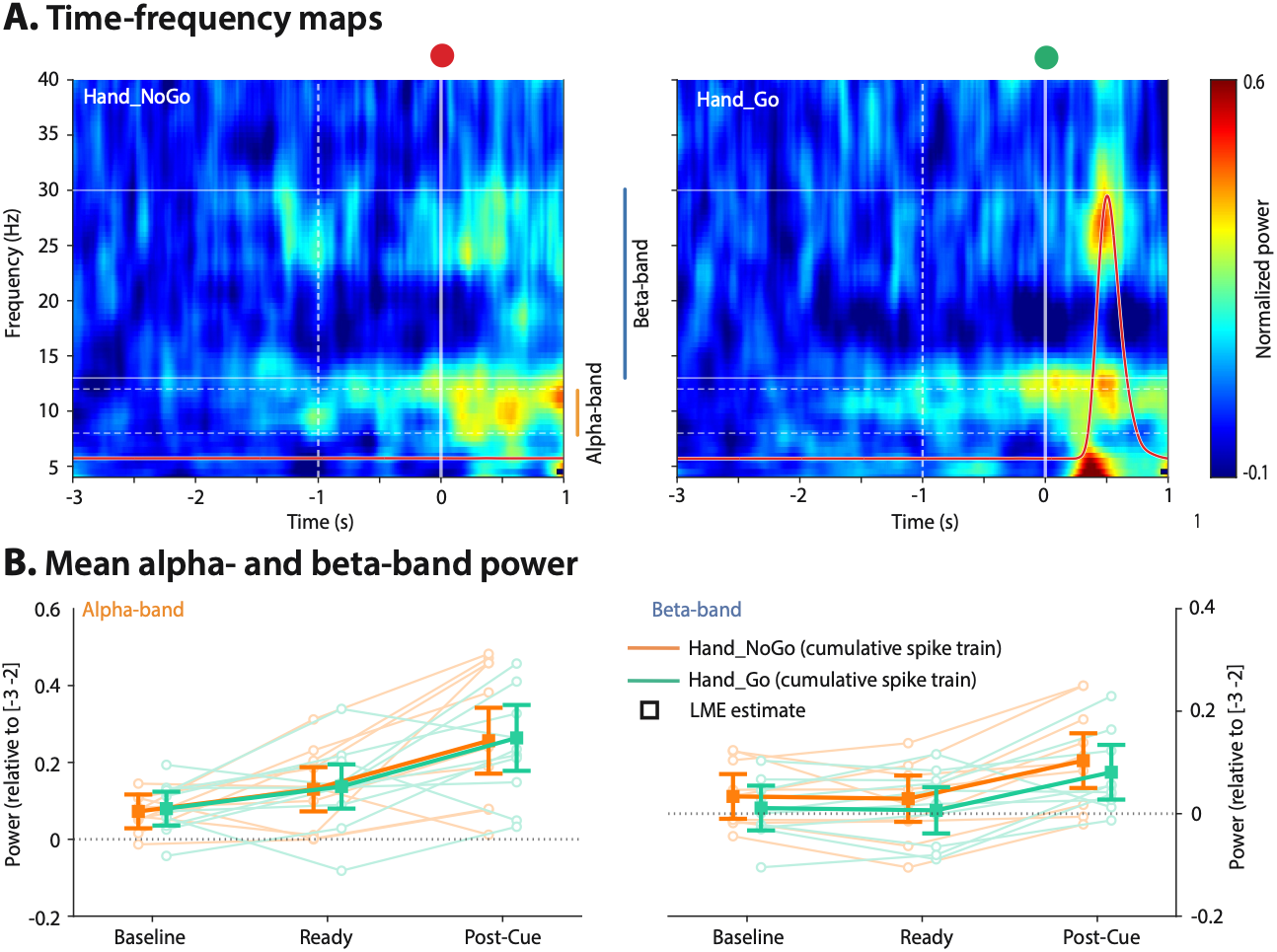
Alpha-and beta-band modulations of cumulative spike train during Hand_NoGo and Hand_Go. In the subset of participants with available motor neuron data in the Hand_Go task (n = 10), we performed complementary analyses comparing alpha-and beta-band cumulative spike train (CST) modulations between conditions. **(A)** Group-averaged time-frequency maps of the CST for Hand_NoGo and Hand_Go. Dashed white vertical lines indicate the Ready cue; solid white vertical lines indicate the imperative cue (red dot: No-Go; green dot: Go). Horizontal dashed white lines mark the alpha-band (8–12 Hz) and beta-band (13–30 Hz) boundaries. For Hand_Go, the mean group force trace (overlaid in red) is shown to indicate the timing of the ballistic response. **(B)** Mean normalized power in the alpha (left) and beta (right) bands for Hand_NoGo and Hand_Go. Thin lines represent individual participants; thick lines and squares represent the linear mixed-effects model estimates.

**Table S1.** Simulation parameters.

| Parameter | Definition | Value |
|---|---|---|
| ***T<sub>sim</sub>*** | Simulation duration (ignoring trimmed edges) | 6 s (fixed) |
| ***dt*** | Integration time step | 0.1 ms (fixed) |
| ***N*** | Number of motor neurons | 40 (fixed) |
| ***V<sub>rest</sub>*** | Resting potential | 0 mV (fixed) |
| ***V<sub>thresh</sub>*** | Firing threshold | 10 mV (fixed) |
| ***t<sub>refractory</sub>*** | Refractory period | 5 ms (fixed) |
| ***D<sub>min</sub>*** | Minimum soma diameter (left truncation of distribution) | 50 µm (fixed) |
| ***D<sub>max</sub>*** | Maximum soma diameter (right truncation of distribution) | 100 µm (fixed) |
| ***D<sub>τ</sub>*** | Scaling factor of the exponential distribution of motor neuron soma diameters | First-stage prior: [0, 10] µm<br>Second-stage: sampled from first-stage posterior |
| ***R<sup>i</sup>*** | Membrane resistance | Size-dependent |
| ***C<sup>i</sup>*** | Membrane capacitance | Size-dependent |
| ***I<sup>i</sup><sub>rheobase</sub>*** | Rheobase | Size-dependent |
| ***τ<sup>i</sup>*** | Membrane time constant (= *R<sup>i</sup>* × *C<sup>i</sup>*). | Size-dependent |
| ***S<sup>i</sup>*** | Input scaling (normalized resistance) | Size-dependent |
| ***V<sub>AHP</sub>*** | Afterhyperpolarization potential | 0 mV (fixed) |
| Δ***g<sub>AHP</sub>*** | Spike-triggered afterhyperpolarization conductance | 10 mS (fixed) |
| ***τ<sup>i</sup><sub>AHP</sub>*** | Afterhyperpolarization time constant | Size-dependent |
| ***I<sub>drive</sub>*** | Baseline net excitatory input (common input offset). | First-stage prior: [35, 45] µA<br>Second-stage: sampled from first-stage posterior |
| ***I<sub>low−freq_f</sub>*** | Low-frequency input low-pass filter cutoff frequency | 5 Hz (fixed) |
| ***I<sub>low−freq_σ</sub>*** | Low-frequency input fluctuation amplitude | First-stage prior: [0.5, 5] µA<br>Second-stage: sampled from first-stage posterior |
| ***I<sub>α_f</sub>*** | Alpha-band input band-pass filter limits | 8 to 13 Hz (fixed) |
| ***I<sub>α_σ</sub>*** | Alpha-band input fluctuation amplitude during baseline window (after envelope modulation) | First-stage prior: [0, 7.5] µA<br>Second-stage: sampled from first-stage posterior |
| ***I<sub>α_envelope_f</sub>*** | Alpha-modulating envelope low-pass filter cutoff frequency | 3 Hz (fixed) |
| ***I<sub>β_f</sub>*** | Beta-band input band-pass filter limits | 13 to 30 Hz (fixed) |
| ***I<sub>β_σ</sub>*** | Beta-band input fluctuation amplitude during baseline window (after envelope modulation) | First-stage prior: [0, 10] µA<br>Second-stage: sampled from first-stage posterior |
| ***I<sub>β_envelope_f</sub>*** | Beta-modulating envelope low-pass filter cutoff frequency | 10 Hz (fixed) |
| ***Burst<sub>t</sub>*** | Center of the burst Gaussian kernel relative to the imperative cue (common to all burst input components) | 0.25 s (fixed) |
| ***Burst<sub>σ</sub>*** | Standard deviation of the burst Gaussian kernel (common to all burst input components) | 100 ms (fixed) |
| ***Trend<sub>t</sub>*** | Start of the linear trend relative to the imperative cue (common to all trend input components) | -1 s (fixed) |
| ***I<sub>drive_burst</sub>*** | Scaling of the Gaussian kernel added to the net excitatory drive | First-stage (disabled): 0 µA<br>Second-stage prior: [-6, 6] µA |
| ***I<sub>drive_trend</sub>*** | Slope of the linear trend added to the net excitatory drive | First-stage (disabled): 0 µA/s<br>Second-stage prior: [-5, 0] µA/s |
| ***I<sub>α_burst</sub>*** | Scaling of the Gaussian kernel added to the alpha-modulating envelope | First-stage (disabled): 0 µA<br>Second-stage prior: [-3, 3] µA |
| ***I<sub>α_trend</sub>*** | Slope of the linear trend added to the alpha-modulating envelope | First-stage (disabled): 0 µA/s<br>Second-stage prior: [-5, 5] µA/s |
| ***I<sub>β_burst</sub>*** | Scaling of the Gaussian kernel added to the beta-modulating envelope | First-stage (disabled): 0 µA<br>Second-stage prior: [-3, 3] µA |
| ***I<sub>β_trend</sub>*** | Slope of the linear trend added to the beta-modulating envelope | First-stage (disabled): 0 µA/s<br>Second-stage prior: [-5, 5] µA/s |
| ***w<sub>common_σ</sub>*** | Target standard deviation of mean-one motor-neuron-specific common input fluctuations scaling weights, sampled from a truncated lognormal distribution | First-stage prior: [0, 2]<br>Second-stage: sampled from first-stage posterior |
| ***w<sub>common_max</sub>*** | Truncation of the lognormal distribution of common input fluctuations scaling weights | 10. (fixed) |
| ***I<sub>independent_f</sub>*** | Independent input frequency range | 0 to 50 Hz (fixed) |
| ***I<sub>independent_σ</sub>*** | Independent input fluctuation amplitude | First-stage prior: [0, 10] µA<br>Second-stage: sampled from first-stage posterior |
| ***w<sub>independent_σ</sub>*** | Target standard deviation of mean-one motor-neuron-specific independent input scaling weights, sampled from a truncated lognormal distribution | First-stage prior: [0, 2]<br>Second-stage: sampled from first-stage posterior |
| ***w<sub>independent_max</sub>*** | Truncation of the lognormal distribution of independent input scaling weights | 10 (fixed) |

